# DAF-16 and SMK-1 contribute to innate immunity during adulthood in *Caenorhabditis elegans*

**DOI:** 10.1101/761312

**Authors:** Daniel R. McHugh, Elena Koumis, Michelle Schlaubitz-Garcia, Safae Bennani, Paul Regan, Prem Patel, Matthew J. Youngman

## Abstract

Aging is accompanied by a progressive decline in immune function termed “immunosenescence”. Deficient surveillance coupled with the impaired function of immune cells compromises host defense in older animals. The dynamic activity of regulatory modules that control immunity appears to underlie age-dependent modifications to the immune system. In the roundworm *Caenorhabditis elegans* levels of the PMK-1 p38 MAP kinase diminish over time, reducing the expression of immune effectors that clear bacterial pathogens. Along with the PMK-1 pathway, innate immunity in *C. elegans* is regulated by the insulin signaling pathway. Here we asked whether DAF-16, a Forkhead box (FOXO) transcription factor whose activity is inhibited by insulin signaling, plays a role in host defense later in life. While in younger worms DAF-16 remains in an inactive state unless stimulated by environmental insults, we found that even in the absence of acute stress the transcriptional activity of DAF-16 increases in an age-dependent manner. Beginning in the reproductive phase of adulthood, DAF-16 upregulates a subset of its transcriptional targets, including genes required to kill ingested microbes. Accordingly, DAF-16 has little to no role in larval immunity, but functions specifically during adulthood to confer resistance to bacterial pathogens. We found that DAF-16-mediated immunity in adults requires SMK-1, a regulatory subunit of the PP4 protein phosphatase complex. Our data suggest that as the function of one branch of the innate immune system of *C. elegans* (PMK-1) declines over time, DAF-16-mediated immunity ramps up to become the predominant means of protecting adults from infection, thus reconfiguring immunity later in life.

## INTRODUCTION

The age-dependent decline in immune function, known as immunosenescence, severely restricts an organism’s ability to defend itself against infectious agents, leading to an enhanced susceptibility to pathogens later in life. In humans and other vertebrate species, the most distinctive features of immunosenescence are those that affect adaptive immunity, and T cells in particular. Aging is accompanied by involution of the thymus, and the T cell pool comes to be dominated by memory cells, leaving relatively little space to be occupied by naïve cells that provide protection from newly encountered pathogens (reviewed in Franceschi *et al.* 2000). Immunosenescence is also characterized by a weakened innate immune response, as revealed by studies of individual types of leukocytes. For example, neutrophils from older animals exhibit reduced chemotaxis (Brubaker *et al.* 2013). In addition, older dendritic cells have a diminished capacity to phagocytose and present antigens, thus reducing T cell priming (Chougnet *et al.* 2015). While many of the hallmarks of immunosenescence have been described, less is understood about potential age-dependent changes at the molecular level in the underlying regulatory pathways that govern host defense. Since cross-talk between pathways facilitating communication between multiple cell types adds to the complexity of this problem, studies to address it at a more fundamental level have benefited from experimental systems such as worms and flies that lack the confounding effects of a bipartite immune system.

Invertebrates retain several components of innate immunity, demonstrating that key elements of immune protection are evolutionarily conserved. Circulating hemocytes with the ability to either sequester, engulf, or poison pathogens are part of the defense arsenal of coelomate metazoans, including the fruitfly, *Drosophila melanogaster* (reviewed in Melillo *et al.* 2018). Other invertebrates, such as the roundworm *Caenorhabditis elegans*, lack true circulating immune cells and instead have an even simpler, more ancient immune system. While the means by which worms detect and distinguish between microbes is still unclear, exposure to pathogens triggers a humoral response involving the production of antimicrobial peptides (AMPs) (Dierking *et al.* 2016). AMPs are predicted to be secreted by intestinal epithelial cells in a manner that is reminiscent of mucosal immunity in other species. The expression of AMPs and other proteins important for responding to acute infection in *C. elegans* is regulated by three evolutionarily conserved signaling pathways: the TGF-ß pathway (Zugasti and Ewbank 2009), the PMK-1 p38 MAPK pathway (Kim *et al.* 2002), and the insulin signaling pathway (Garsin *et al.* 2003), each of which has been shown to modulate the function of immune cells in mammals (Hayakawa *et al.* 2017; Fischer *et al.* 2017; Batlle and Massagué 2019).

Over the last 20 years, several studies have shown that, similar to vertebrates, the ability of flies and worms to resist infection diminishes with age, with older animals dying from acute infection more rapidly than young animals (Tan *et al.* 1999; Laws *et al.* 2004; Ramsden *et al.* 2008; Youngman *et al.* 2011). This suggests that the factors responsible for eroding and reshaping immunity over time may be shared among animals that experience aging, regardless of the complexity of their means of host defense. As it has been described thus far, immunosenescence in *C. elegans* seems to be brought on by three age-related changes that directly infringe upon the expression of AMPs. The first is a collapse in proteostasis beginning in early adulthood (Ben-Zvi *et al.* 2009 p.; Taylor and Dillin 2013) that is coupled with accumulation of vitellogenin in the intestine (Herndon *et al.* 2002; Seah *et al.* 2016). Experimental conditions to mimic these conditions using transgenic animals caused them to be more susceptible to *P. aeruginosa* infection, and mutations in VIT-2 that cause it to build up in the intestine block the expression of genes encoding immune effectors. This leads to a model in which the protein instability later in life triggers the repression of genes encoding secreted proteins including those that function in immunity, yet whether this scenario plays out during normal aging has not been directly tested (Singh and Aballay 2017). A second contributor to immunosenescence in worms is a decline in the activity of the SKN-1 transcription factor over time (Papp *et al.* 2012). In addition to its well-characterized role in oxidative stress resistance (An and Blackwell 2003) SKN-1 is also required for resistance to *P. aeruginosa* infection in both larvae and adults. A third factor in the age-dependent decline in immunity in *C. elegans* is a progressive decrease in the levels of the PMK-1 protein during aging that enhances the susceptibility of older worms to bacterial infection (Youngman *et al.* 2011). Without youthful levels of PMK-1, the worm is left with one less branch of its innate immune system as the expression levels of genes encoding immune effectors regulated by the PMK-1/p38 MAPK pathway decreases. Whether the activity of the two other major pathways that modulate immunity in *C. elegans* also declines over time is not known. Considering its vital roles in both stress resistance and lifespan determination, we hypothesized that the insulin signaling pathway plays an important role in protecting older worms from infection.

Across evolutionarily diverse animal phyla, the insulin and insulin-like growth factor signaling pathway (IIS) plays the dual role of regulating lifespan and integrating environmental cues to regulate growth and development (Kenyon *et al.* 1993; Tamemoto *et al.* 1994; Kimura *et al.* 1997; Tatar *et al.* 2001; Yuan *et al.* 2009). Two of the key components of the IIS pathway are a transmembrane receptor for insulin/insulin-like ligands and a transcription factor of the Forkhead Box family (FOXO) whose transcriptional activity is regulated by a cascade of kinase-mediated phosphorylation reactions (reviewed in Lapierre and Hansen 2012). In the default status of this pathway when the insulin receptor is bound by a ligand, the ensuing phosphorylation cascade culminates in an inhibitory phosphate group being added to the FOXO transcription factor, preventing its translocation to the nucleus to regulate the expression of its target genes. This inhibition may be overcome, however, when an animal encounters an acute stress. For example, in *C. elegans* under circumstances of starvation, irradiation, or other insults, the FOXO transcription factor is de-repressed and upregulates the expression of genes important for neutralizing the stress or counteracting its effects (Henderson and Johnson 2001; Essers *et al.* 2005). Manipulations to artificially activate the FOXO transcription factor (for example, by overexpressing FOXO or disrupting the function of the insulin receptor) dramatically extend the worm’s lifespan. Such experiments suggest that variability in the degree of FOXO activity during aging could help to explain how some individuals live longer and in better health than others. Indeed, in synchronized aging cohorts of *Caenorhabditis elegans*, animal-to-animal differences in FOXO activity have been observed, with greater levels of activity positively correlating with longer lifespan (Sánchez-Blanco and Kim 2011).

In *C. elegans* DAF-16 is the orthologue of mammalian FOXO3a, and its transcriptional activity is antagonized by the IIS when the DAF-2 receptor is occupied by insulin-like peptides. Much of what is understood about the function of DAF-16 has been revealed through studies of *daf-2* mutants in which the constitutive activation of DAF-16 more than doubles the worms’ lifespan (Kenyon *et al.* 1993). One of the key elements that underlies the extended lifespan of *daf-2* mutants is their remarkable resilience when challenged with environmental insults. *daf-2* mutants are resistant to an astounding array of stressors including hypoxia (Jiang *et al.* 2011), anoxia (Garcia *et al.* 2015), oxidative stress, thermal stress, and ultraviolet light (Murakami and Johnson 1996; Wolff *et al.* 2006) as well as fungal and bacterial pathogens (Garsin *et al.* 2003; Wolff *et al.* 2006; Kerry *et al.* 2006). In each case the enhanced resistance of *daf-2* animals, as demonstrated by their ability to outlive wild type animals subjected to the same stressors, is completely dependent upon *daf-16*, implying that DAF-16 modulates the transcriptional response to environmental stress. Supporting this possibility, when animals are exposed to stressful conditions, DAF-16 translocates from the cytosol to the nucleus. For example, in both L2 larvae and in mature adult worms, the oxidative stress inducing compound juglone results in nuclear accumulation of DAF-16::GFP (Henderson and Johnson 2001). Starvation, heat, and ultraviolet irradiation are other stimuli that drive DAF-16::GFP into the nucleus in *C. elegans* (Lin *et al.* 2001; Henderson and Johnson 2001; Liang *et al.* 2006; Evans *et al.* 2008; Singh and Aballay 2009; Mueller *et al.* 2014). This phenomenon is evolutionarily conserved, as oxidative stress also induces the nuclear translocation of FOXO3 and FOXO4 in mammalian cells (Brunet *et al.* 2004; Kobayashi *et al.* 2005). Multiple transcriptomic studies using a variety of platforms have demonstrated that once inside the nucleus, DAF-16 upregulates genes whose products function to directly neutralize environmental threats or to repair the damage caused by them (McElwee *et al.* 2003; Lee *et al.* 2003; Murphy *et al.* 2003; Tepper *et al.* 2013; Chen *et al.* 2015; Li *et al.* 2019). These include antimicrobial peptides, superoxide dismutases, and chaperones. The basis for the long life of *daf-2* mutants, therefore, appears to lie at least in part in the enhanced expression of DAF-16 transcriptional targets that buffer animals against environmental perturbations and that maintain cellular health.

As DAF-16 is the foremost contributor to stress resistance in *daf-2* animals, it would stand to reason that, in a corresponding manner, *daf-16* mutants themselves would exhibit sensitivity to the same stresses to which *daf-2* mutants are resistant. For the most part, this is indeed the case. When DAF-16 function is compromised by either loss-of-function mutations or by RNAi, animals die more rapidly than wild type controls when subjected to hypoxic conditions (Jiang *et al.* 2011), paraquat and hydrogen peroxide (Nemoto-Sasaki and Kasai 2009), high temperature (Li *et al.* 2008), and heavy metals (Chu *et al.* 2005; Li *et al.* 2008). However, in the absence of systemic DAF-16, larval stage worms do not seem to be more susceptible to infection by opportunistic human pathogenic bacteria (Garsin *et al.* 2003; Liang *et al.* 2006; Evans *et al.* 2008) or fungi (Kerry *et al.* 2006). These observations suggest that although it is important for the defense against other stresses, DAF-16 does not protect younger animals from infection. Curiously, when *C. elegans* are infected as Day 2 adults after transitioning from the L4 stage to reproductive maturity in the presence of RNAi targeting *daf-16* they then succumb to infection with *Pseudomonas aeruginosa* more rapidly than untreated control animals (Twumasi-Boateng *et al.* 2012). This raises the intriguing possibility of an age-dependent role for DAF-16 in innate immunity, but a systematic analysis of the function of DAF-16 during aging to has not yet been performed.

We wondered whether the erosion of PMK-1-mediated immunity during aging may be counterbalanced by the function of DAF-16. To test this possibility, we took a primarily genetic approach to functionally characterize DAF-16 over time, with a particular emphasis on examining its role in post-reproductive adults. During aging in wild type animals maintained under standard (non-stressed) laboratory conditions the transcriptional activity of DAF-16 increases in an age-dependent manner, beginning in early adulthood. Analysis of genome-wide changes in expression during aging revealed that in adult *C. elegans* some DAF-16 targets are upregulated while others are downregulated, suggesting that DAF-16 acts as both a transcriptional activator and a repressor in those animals. An important functional consequence of activating DAF-16 during aging is immune protection, and we find that DAF-16 functions specifically during adulthood to protect aging animals from bacterial infection. In its role in innate immunity DAF-16 appears to cooperate with SMK-1, a regulatory component of the protein phosphatase 4 (PP4) complex. Our data thus establish that DAF-16 is functional during normal healthy aging in wild type animals. Further, they imply that the activation of DAF-16 in adults may be part of a compensatory mechanism to preserve organismal health in response to the flagging activity of other stress response pathways.

## MATERIALS AND METHODS

### *C. elegans* strains and maintenance

Worms were maintained using standard techniques as previously described (Brenner 1974). The *C. elegans* strains used in this study are as follows: Bristol wildtype N2, *pmk-1(km25)*, *daf-16(mgDf47)*, *pmk-1(km25);daf-16 (mgDf47)*, *daf-2(e1368)*, *daf-2(e1370)*, TJ356 *zIs356* [*daf-16p::daf-16a/b::GFP + rol-6(su1006)*], and VIL001 *mjyIs001* [*plys-7::GFP*].

To generate strain VIL001, stable transgenic animals harboring an extrachromosomal *lys-7* promoter::gfp fusion construct (Alper *et al.* 2007) were subjected to gamma irradiation to yield a chromosomal integration of *plys-7::gfp*. F_2_ segregants of the irradiated P_0_ animals yielding 100% GFP-expressing progeny were selected for further analysis. One of these lines was backcrossed to the N2 wildtype strain seven times and then designated VIL001.

### Generation of age-matched cohorts of *C. elegans*

Synchronized populations of worms were obtained by sodium hypochlorite treatment of gravid adult animals to harvest eggs followed by hatching L1 larvae in M9 buffer without nutritional supplementation for 16-20h at 22°C. Approximately 2000 L1 larvae were plated on to NGM media seeded with either *E. coli* OP50 *or E. coli* HT115 strains harboring RNAi constructs, depending on the experiment. To age animals to the sixth day of adulthood, worms were allowed to develop to the L4 larval stage before transferring them to new NGM plates containing 25 ug/mL 5-fluorodeoxyuridine (FUdR) and seeded with *E. coli*.

### RNAi treatment

*E. coli* HT115 strains carrying plasmids encoding dsRNA were thawed from glycerol stocks stored at −80°C and grown overnight at 37°C on LB plates supplemented with ampicillin and tetracycline. Single colonies were used to inoculate 200 mL of LB + ampicillin and incubated with shaking overnight at 37° C. To concentrate cells, cultures were centrifuged for 10 minutes at 3000x*g* and the cell pellet was resuspended in 25 mL of fresh LB supplemented with ampicillin. Approximately 1 mL of the concentrated cell cultures was used to seed two sets of NGM plates: one containing carbenicillin, and 2M IPTG (referred to as “RNAi plates”) and a second set containing carbenicillin, 2M IPTG, and 25ug/mL FUdR; referred to as “RNAi + FUdR plates”). Seeded plates were incubated in the dark at 22°C for at least 3 days before worms were added to them for knockdown experiments.

Synchronized populations of worms were subjected to RNAi treatments at one of two points during aging, depending on the experiment. For experiments involving a brief pulse of RNAi to knockdown *daf-16*, L1 larvae harvested from hypochlorite treatment of gravid adult hermaphrodites were maintained on NGM plates with *E. coli* OP50 as the food source at 20°C as described above until the fourth day of adulthood. At 48 hours prior to initiating infection with *P. aeruginosa*, these adult worms were transferred to RNAi +FUdR plates seeded with the *daf-16* RNAi clone (or L4440 vector control clones) and incubated at 20°C. For all other RNAi experiments, after hatching overnight in M9 buffer L1 larvae harvested from hypochlorite treatment of gravid adult hermaphrodites were introduced to concentrated *E. coli* RNAi clones seeded on to RNAi plates and maintained on those plates at 20°C. Once these animals reached the L4 stage, they were transferred by chunking to RNAi + FUdR plates where they were allowed to move away from the agar chunk before it was removed from the plate. Worms were aged on these plates at 20°C until the sixth day of adulthood when they were harvested or used for other assays.

### Fluorescence microscopy

To analyze the expression of the *plys-7::GFP* reporter during aging, worms maintained on NGM plates seeded with OP50 were mounted onto agarose pads (10% agarose in M9 buffer) seated on glass slides and immobilized in a slurry of 0.1 μm polystyrene beads (Polysciences, Warrington, PA). Samples were examined by compound fluorescence microscopy on a Zeiss Axioskop (Zeiss, Oberkochen, Germany), and images were captured using a Zeiss digital camera. To study the effect of different RNAi treatments on the expression of *lys-7::GFP*, worms on RNAi or RNAi + FUdR plates were inspected under a fluorescence dissecting microscope (Zeiss) equipped with a Zeiss digital camera. For all imaging studies, camera settings including exposure time were held constant for all samples.

### *Pseudomonas aeruginosa* infection

Infections of larval stage and adult *C. elegans* with *P. aeruginosa* were carried out as described (Youngman *et al.* 2011). Briefly, approximately 100 worms of the indicated ages were transferred from plates containing *E. coli* OP50 or *E. coli* HT115 (for RNAi treatments) to plates seeded with *P. aeruginosa* strain PA14 and incubated at 25°C. The survival of worms on the *P. aeruginosa* plates was assessed and dead worms were removed every 12-24 hours until all of the worms in the assay had died. Each assay was replicated at least twice.

### Calculation of median survival times and statistical analysis

To calculate the median survival time (LT_50_) of worms infected with *P. aeruginosa*, the fraction of animals alive at each time point during an assay was first plotted as a function of time in Excel, accounting for animals that were inadvertently overlooked or that escaped between time points. These data were then imported into SigmaPlot (Systat Software, San Jose, CA) and a three parameter sigmoidal curve was fit according to the general equation y=a/(1=e^-(x-x0)/b^). This equation was used to determine the point at which 50% of the animals in the assay had died. The average fold difference between the LT_50_ of mutant strains or experimental RNAi treatments and control animals was calculated, and the statistical significance of that difference was assessed using a two-tailed Student’s t-test.

### RNA isolation

Approximately 5,000-10,000 worms were washed off of NGM plates in M9 buffer and allowed to settle to the bottom of a 15 mL conical tube by gravity. After removing the supernatant the worms were washed with 10 mL of fresh M9 and again allowed to settle to the bottom of the tube. All but 1-2 mL of buffer were removed and used to resuspend the worms before transferring them to cryovials where more buffer was removed, leaving behind sufficient volume to cover the worm pellet. After adding 300 μL of Trizol reagent (ThermoFisher Scientific, Waltham, MA), the worms were vortexed in a series of 30 second intervals over a period of 5 minutes interspersed by brief rest periods and then transferred to −80°C for storage. To prepare total RNA from worm pellets, frozen worms in Trizol were thawed at room temperature and then briefly vortexed before pelleting in a microcentrifuge at 16,000x*g* for 5 min to remove worm carcasses and debris. Following a phenol/chloroform extraction, total RNA was precipitated in isopropanol, washed in 70% ethanol, and resuspended in RNase-free H_2_O. For RNAseq experiments, total RNA was treated with DNase and reisolated over a column using the RNeasy kit (Qiagen, Germantown, MD). The concentration of RNA was measured by either Nanodrop (ThermoFisher Scientific, Waltham, MA) or by Qubit fluorimetry (Invitrogen, Carlsbad, CA).

### mRNAseq analysis

Prior to RNAseq analysis, total RNA harvested from N2 L4 or Day 6 adult *C. elegans* fed the RNAi vector control *E. coli* strain HT115 was first scrutinized for evidence of degradation and the presence of robust peaks corresponding to ribosomal RNA (rRNA) using a Bioanalyzer (Agilent Technologies, Santa Clara, CA) and agarose gel electrophoresis. rRNA was then depleted using Ribo-Zero (Illumina, San Diego, CA), and cDNA libraries were prepared by strand-specific cDNA synthesis. After determining the median library insert size and total yield, samples were sequenced using Illumina HiSeq 2500 1x 125 single read technology.

### Bioinformatics methods

Bioinformatic analysis was carried out by iGenBio (Chicago, IL) using proprietary ERGO 2.0 software. Briefly, fastqc sequence files were checked for ribosomal depletion, contamination, leftover adapter sequences, and other issues, and the samples were trimmed. All samples were aligned against the reference genome (Wormbase version WS254) using TopHat2. The alignments were then run with featureCounts obtaining a table of counts for each feature in the reference genome. The counts were then imported into R, and DESeq2 was run on the samples. Before each analysis in ERGO the data is normalized using edgeR’s calcNormFactors, then transformed using limma’s voom. Ranked lists of genes differentially expressed between Day 6 adults and L4 larvae were generated with corresponding *p* and *q* values for each gene.

To identify genes with statistically significant changes in expression between the L4 larval stage and Day 6 of adulthood, we eliminated from our list of differentially expressed genes any gene that had a *q* value of greater than 0.05. For a gene to be considered as either up- or downregulated, the difference in its expression levels between the L4 larval stage and Day 6 of adulthood had to be greater than 2-fold, but for this study we considered only those genes whose expression changed by 5-fold for more. To generate a complete catalog DAF-16 transcriptional targets, we created a composite list of non-duplicate entries for genes that are both positively- and negatively regulated by DAF-16 reported in three different studies (Tepper *et al.* 2013; Chen *et al.* 2015; Li *et al.* 2019). The catalog was then compared to our filtered comprehensive lists of genes differentially regulated during aging. Functional information regarding the products of genes differentially regulated during aging or of DAF-16 transcriptional targets was obtained through analysis of gene lists using Gene Ontologies through the Panther database (pantherdb.org) and were focused on GO biological process and child cellular process annotations.

### qRT-PCR

For analysis of gene expression levels by qRT-PCR, cDNA was reverse transcribed from 1 μg samples of total RNA using Retroscript (ThermoFisher Scientific, Waltham, MA). Diluted cDNA was used as a template in PCR reactions designed to amplify sequences from *lys-7*, *sod-3*, and *mtl-1* transcripts with *tba-1* as the reference using primer pairs described previously (Troemel *et al.* 2006). Sybr Green (Roche, New York, NY) was used to detect the products of the PCR reactions, and fluorescence was quantified using an Eppendorf Master Cycler (Eppendorf, Hauppauge, NY). Relative gene expression levels were calculated using the ΔΔCt method (Livak and Schmittgen 2001). This analysis was performed on a total of three biological replicates, representing three independently generated cohorts of aging worms.

### Data availability

RNAseq data will be made available on the GEO database (ncbi.nlm.nih/geo). Files uploaded as supplemental material to figshare are described below. Figure S1 shows the results of a second replicate of the experiment investigating the function of individual isoforms of DAF-16 (Fig. 3). Figures S2 and S3 depict pie charts of overrepresented GO biological process and cellular process terms in genes that are up- and downregulated at Day 6 of adulthood, respectively. Figures S4 and S5 show pie charts of overrepresented GO biological process and cellular process terms in DAF-16 targets that are up- and downregulated at Day 6 of adulthood, respectively. Figure S6 shows a second replicate of the *P. aeruginosa* infection assay involving *daf-2* animals (Fig. 5). It also shows survival curves of animals treated with RNAi targeting *daf-2* and then infected with *P. aeruginosa* at the L4 larval stage and Day 6 of adulthood. Table S1 lists genomewide changes in expression between the L4 larval stage and Day 6 of adulthood. Tables S2 and S3 list overrepresented GO biological process terms associated with genes up- and downregulated at Day 6, respectively. Table S4 is a catalog of DAF-16 targets made by compiling lists of DAF-16 targets identified in previous reports (Tepper *et al.* 2013; Chen *et al.* 2015; Li *et al.* 2019). Table S5 lists DAF-16 target genes that are differentially expressed between the L4 larval stage and Day 6 of adulthood. Tables S6 and S7 list overrepresented GO biological process terms associated with DAF-16 targets that are up- and downregulated at Day 6, respectively.

## RESULTS

### Comparison of PMK-1 and DAF-16 function in innate immunity

Although they regulate separate groups of genes, both *pmk-1* and *daf-16* are necessary for lifespan extension of *daf-2(e1370)* mutants, suggesting that there may be some degree of functional overlap between the two gene products even though they are members of distinct genetic pathways (Troemel *et al.* 2006). PMK-1 and DAF-16 also confer enhanced resistance to pathogens in larval stage *daf-2(e1370)* animals, indicating that each may contribute to longevity through at least partially redundant functions in innate immunity. Whether the two proteins operate in a similar manner in the context of aging in wild type animals is not known. To compare the functions of DAF-16 and PMK-1 in innate immunity during aging, we examined the survival of *pmk-1(km25)*, *daf-16(mgDf47)* and *pmk-1(km25);daf-16(mgDf47)* mutants upon infection with the human opportunistic bacterial pathogen *Pseudomonas aeruginosa* at the fourth larval stage (L4) and at Day 6 of adulthood (D6; six days after the transition to L4). Consistent with previous reports, when bacterial infection was initiated at the L4 larval stage, *pmk-1(km25)* mutants died more rapidly from the infection than did wild type animals, confirming a role for PMK-1 in the innate immunity of juvenile worms (Fig. 1A). The *daf-16(mgDf47)* loss-of-function mutation, on the other hand, did not affect the ability of L4 worms to resist bacterial infection. In addition, *daf-16(mgDf47)* did not further enhance the susceptibility of animals bearing a lesion in the *pmk-1* locus, as *pmk-1(km25);daf-16(mgDf47)* double mutants died from infection at the same rate as *pmk-1(km25)* mutants (Fig 1A). These data suggest that DAF-16 plays little to no role in innate immunity in L4 larvae and that host defense is mediated primarily by PMK-1 at that stage. When infection was initiated at Day 6 of adulthood we observed a different pattern in the survival of the mutants (Fig. 1B). While *pmk-1(km25)* mutants still died faster from the infection than wild type animals, the median lifespan of infected *daf-16(mgDf47)* mutants was also reduced. In addition, the *pmk-1(km25);daf-16(mgDf47)* double mutant displayed an additive phenotype, with the shortest median lifespan of all the strains. These results suggest that by Day 6 of adulthood a second arm of innate immunity mediated by DAF-16 has become activated and that it functions in parallel to PMK-1.

**Figure 1.**
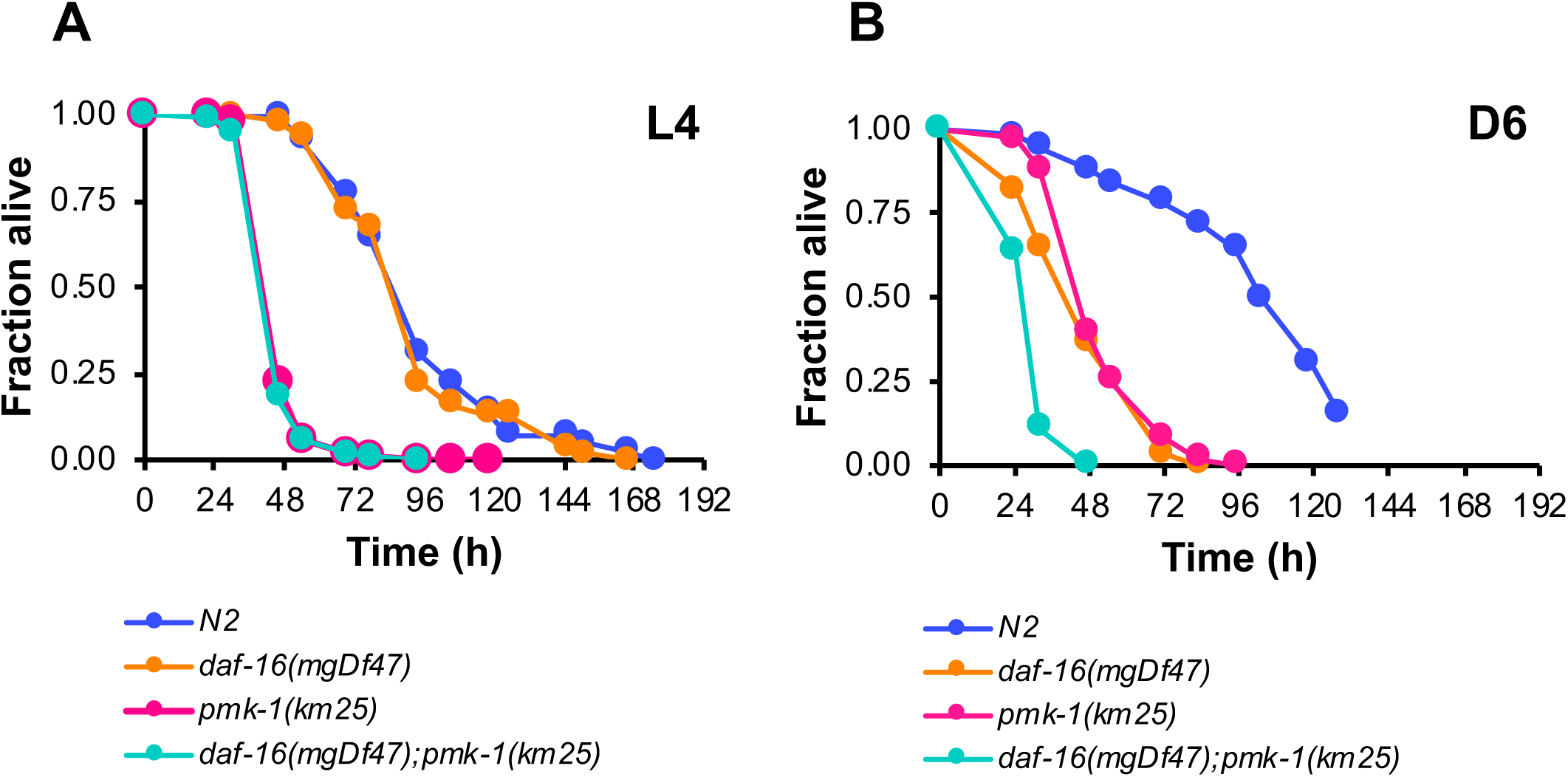
DAF-16 likely functions in parallel to PMK-1 to modulate innate immunity during aging. (A) Survival of wild type strain N2 (blue), and mutant strains *daf-16(mgDf47)* (orange), *pmk-1(km25)* (pink), and *daf-16(mgDf47);pmk-1(km25)* (green) transferred from *E. coli* OP50 to *P. aeruginosa* PA14 at L4. (B) Survival of the same strains as in (A) upon initiating infection with *P. aeruginosa* at Day 6 of adulthood. In both panels, the fraction of worms alive is plotted as a function of time.

### Systematic functional analysis of DAF-16 in innate immunity during aging

To systematically address the potential role for DAF-16 in innate immunity in adult worms, we asked about its functional requirement in host defense over time by challenging *daf-16(mgDf47)* animals with *P. aeruginosa* infection at four different points during their lifespan. In parallel, age-matched populations of wild type N2 animals were also exposed to pathogen, and their survival was compared to that of the infected *daf-16(mgDf47)* mutants. While the absence of functional DAF-16 had little to no effect on the ability of L4 worms to resist infection (Fig. 2A), the age-dependent increase in susceptibility to bacterial pathogens observed in wild type worms was exacerbated in *daf-16(mgDf47)* mutants. This was evident at Day 3 of adulthood (Fig 2B). Day 3 adult N2 worms were no more susceptible to bacterial infection than wild type L4 larvae. However, the average median survival time (LT_50_) of Day 3 *daf-16(mgDf47)* was 1.4-fold less than the LT_50_ of wildtype control animals (p<0.001, Table 1). This difference in survival was more pronounced when worms were challenged with *P. aeruginosa* at Day 6 of adulthood (Fig. 2C). The conditions used in our studies successfully recapitulated previous reports of declining immunity in adult worms, manifested as increased susceptibility to bacterial infection over time (Laws *et al.* 2004; Youngman *et al.* 2011). The absence of DAF-16 caused Day 6 adults to die even more rapidly from infection. The median survival time of Day 6 *daf-16(mgDf47)* worms challenged with *P. aeruginosa* was less than half of the LT_50_ of N2 wildtype animals (p= 0.0019, Table 1). The loss-of-function mutation in *daf-16* still had an effect when Day 9 worms were infected with *P. aeruginosa*, resulting in a 1.4-fold reduction in LT_50_ of *daf-16(mgDf47)* mutants as compared to control worms (p= 0.0322, Fig. 2D, Table 1). Our observations suggest that despite playing only a minor role, if any, in innate immunity in larval stage worms, DAF-16 becomes increasingly important for host defense during aging.

**Figure 2.**
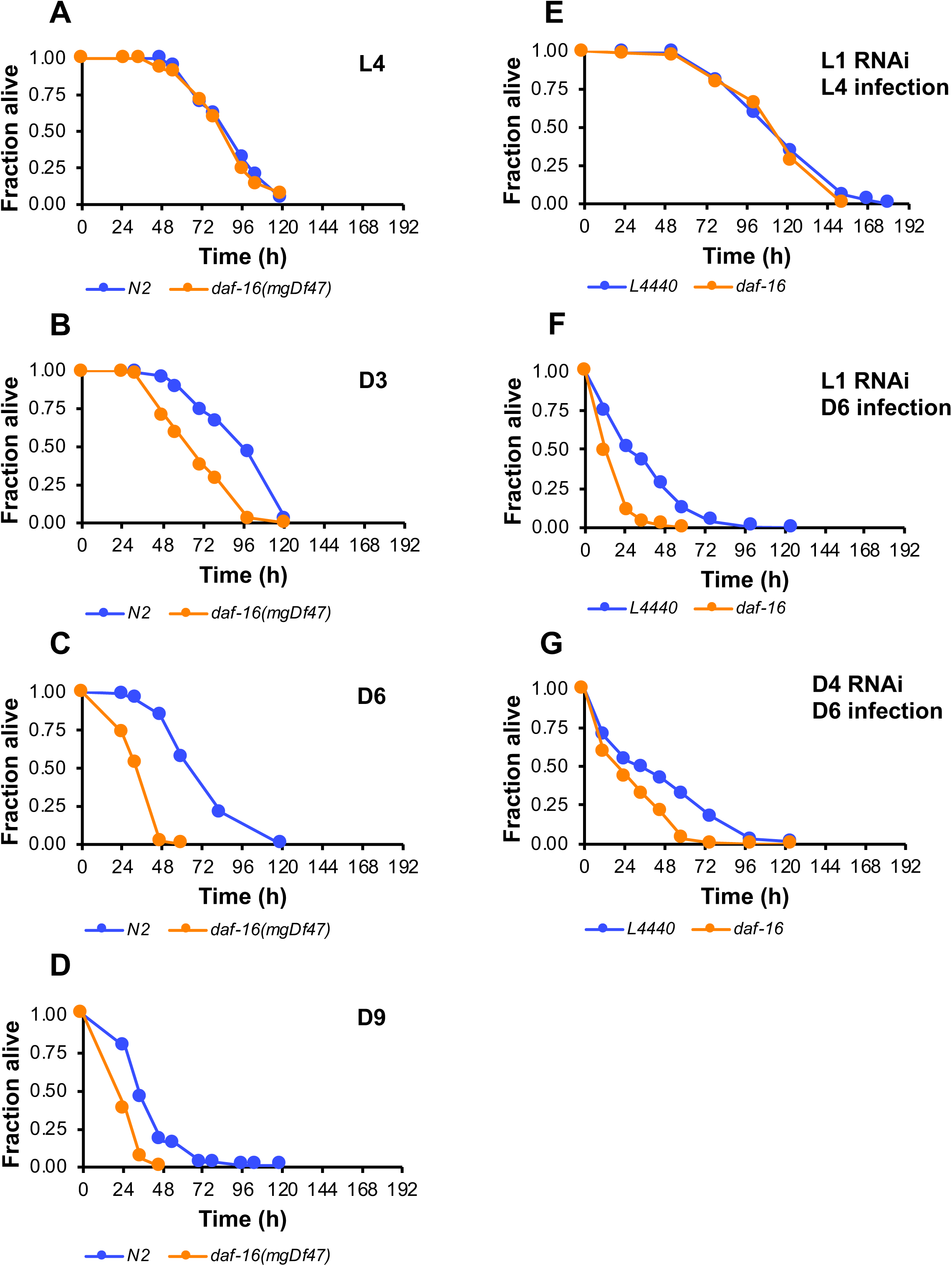
DAF-16 confers resistance to bacterial pathogens during adulthood. (A-D) Wild type strain N2 (blue) or *daf-16(mgDf47)* (orange) mutants were maintained on *E. coli* OP50 and then challenged with *P. aeruginosa* PA14 at the L4 larval stage (A) or at the indicated day of adulthood (B-D). (E-G) RNAi was used to test the timing requirement of DAF-16 in innate immunity. After initiating knockdown of *daf-16* by feeding-based RNAi (orange) at the L1 larval stage (E and F) or at Day 4 of adulthood (G), worms were infected with *P. aeruginosa* PA14 as L4 larvae (E) or at Day 6 of adulthood (F and G). L4440 (blue) is an empty RNAi vector control. In each case, the fraction of worms alive is plotted as a function of time.

**Table 1.** Relative median lifespans for larval and adult *daf-16* mutants or animals treated with RNAi targeting *daf-16* or *smk-1* upon infection with *P. aeruginosa*.

| Age<sup>a</sup> | Genotype/condition | Average relative median lifespan (+/- SEM)<sup>b</sup> | p-value |
| --- | --- | --- | --- |
| L4 | *daf-16(mgDf47)* | 0.921 ± 0.0208 | 0.00245 |
| D3 | *daf-16(mgDf47)* | 0.707 ± 0.0215 | <0.001 |
| D6 | *daf-16(mgDf47)* | 0.445 ± 0.106 | 0.0019 |
| D9 | *daf-16(mgDf47)* | 0.713 ± 0.0529 | 0.0322 |
| L4 | *daf-16(mu86)* | 0.891 ± 0.360 | 0.00232 |
| D3 | *daf-16(mu86)* | 0.563 ± 0.0148 | <0.001 |
| D6 | *daf-16(mu86)* | 0.511 ± 0.308 | ns |
| L4 | *daf-16* total RNAi | 0.935 ± 0.0112 | 0.0043 |
| L4 | *daf-16a* RNAi | 0.984 ± 0.0164 | ns |
| L4 | *daf-16b* RNAi | 1.014 ± 0.0511 | ns |
| L4 | *daf-16af* RNAi | 0.955 ± 0.0349 | ns |
| L4 | *daf-16abf* RNAi | 0.936 ± 0.0309 | ns |
| L4 | *daf-16df* RNAi | 0.983 ± 0.0301 | ns |
| D6 | *daf-16* total RNAi | 0.440 ± 0.0351 | <0.001 |
| D6 | *daf-16a* RNAi | 0.494 ± 0.0921 | 0.00535 |
| D6 | *daf-16b* RNAi | 0.612 ± 0.0782 | 0.00771 |
| D6 | *daf-16af* RNAi | 0.462 ± 0.0711 | 0.00164 |
| D6 | *daf-16abf* RNAi | 0.406 ± 0.0244 | <0.001 |
| D6 | *daf-16df* RNAi | 0.805 ± 0.170 | ns |
| L4 | *smk-1* RNAi | 0.888 ± 0.0391 | 0.0168 |
| D6 | *smk-1* RNAi | 0.511 ± 0.0428 | <0.001 |

<sup>a</sup>Age of animals at the time that *P. aeruginosa* infection was initiated.
<sup>b</sup>The average median lifespan (LT<sub>50</sub>) of the indicated animals is reported as a fraction of the average median lifespan of age-matched controls (N2 or worms treated with the empty RNAi vector L4440).

### Timing requirement for DAF-16 in adult innate immunity

Since loss of *daf-16* is associated with precocious early aging or “progeric” phenotypes (Samuelson *et al.* 2007) our observation that adult *daf-16(mgDf47)* mutants are more susceptible to bacterial infection than larval stage *daf-16(mgDf47)* animals raises the possibility that instead of playing a direct role in immunity during adulthood, DAF-16 is required throughout the entirety of the worm’s life to protect against premature frailty that would cause adult animals to be especially vulnerable to attack by pathogens. To investigate whether this is true, we asked whether a short pulse of RNAi to knock down *daf-16* just before challenging Day 6 adult worms with *P. aeruginosa* would affect their ability to resist the infection to the same extent as a sustained RNAi regimen targeting *daf-16* from the L1 larval phase up until the time that the infection was initiated at the sixth day of adulthood. We reasoned that if DAF-16 contributes to innate immunity in adult worms by principally functioning during adulthood, then waiting until Day 4 of adulthood to inhibit *daf-16* expression should be sufficient to cause enhanced susceptibility to pathogen upon infecting worms RNAi-treated worms with *P. aeruginosa* at Day 6. On the other hand, if DAF-16 somehow exerts its effect on immunity by acting earlier in life then it should be necessary to begin knocking down *daf-16* while worms are still developing to affect their ability to resist bacterial infections as adults. We found that the brief pulse of RNAi targeting *daf-16* at Day 4 had the same effect on the pathogen susceptibility of Day 6 adults as knocking down *daf-16* beginning at the first larval stage (Fig. 2F and G). Our data indicate that DAF-16-mediated immunity in adult animals is a consequence of the specific function of DAF-16 during adulthood and not before. This is agrees with previous observations indicating that DAF-16 is dispensable in larvae for its role in lifespan determination (Dillin *et al.* 2002).

### Functional analysis of specific isoforms of *daf-16* in adult *C. elegans*

There are twelve different isoforms of *daf-16*, but only a few have been functionally characterized. The two that appear to be especially relevant for stress resistance and aging are *daf-16a* and *daf-16d/f*, but there is a lack of consensus regarding which specific isoform is in fact the predominant lifespan determinant (Kwon *et al.* 2010; Bansal *et al.* 2014; Chen *et al.* 2015). Both isoforms are upregulated during aging after the L4 stage with a much more dramatic increase in expression of *daf-16d/f* as compared to *daf-16a* (Bansal *et al.* 2014). Since the timing of the increase in *daf-16a* and *daf-16d/f* expression generally corresponds to the period when our data indicate that DAF-16 becomes activated during aging, we asked whether either of these isoforms contributes to innate immunity in adult animals. For the sake of completeness, we also included the *daf-16b* isoform as part of our analysis, even though it has been implicated to play only minor roles in dauer formation and longevity (Lee *et al.* 2001; Kwon *et al.* 2010).

To determine whether a specific isoform of *daf-16* is responsible for mediating innate immunity in adult worms, a series of RNAi constructs created by the Tissenbaum lab (Kwon *et al.* 2010) was used to target individual *daf-16* isoforms (*a*, *b*, and *d/f*) or combinations of isoforms (*af*, and *abf*) in age-matched cohorts of wild type N2 worms beginning at the L1 stage and continuing until they were challenged with *P. aeruginosa* as L4 larvae or as Day 6 adults. The survival of infected worms was compared to that of an isogenic age-matched control group that had been subjected to RNAi designed to reduce the expression of all *daf-16* isoforms (referred to as “*daf-16* total”). Regardless of which isoforms were targeted, RNAi knockdown of *daf-16* expression did not affect the resistance of L4 worms to the bacterial pathogen (Fig 3A). When the infection was initiated at Day 6, however, all RNAi treatments had at least some impact on the worms’ survival. Similar to what was observed in other experiments, knockdown of all *daf-16* isoforms by daf-16 total RNAi reduced the LT50 od Day 6 adults by almost 2.3-fold (p<0.001, Fig. 3B and C). Knocking down the d/f isoform of DAF-16 did not cause a statistically significant reduction in the LT50 of Day 6 animals, but RNAi targeting *daf-16b* did. Notably, only those RNAi treatments that included knockdown of the *daf-16a* isoform were sufficient to recapitulate the susceptibility phenotype produced by the *daf-16* total construct. RNAi targeting the *daf-16a* isoform alone or in combination with the *f* isoform or with both *daf-16b* and *daf-16f* together resulted in a reduction in the LT_50_ of *P. aeruginosa*-infected Day 6 animals that approximated what was observed when the expression of all *daf-16* isoforms was inhibited by RNAi (Fig. 3B and C, Fig. S1). These results suggest that multiple *daf-16* isoforms may contribute to host defense in older animals, with *daf-16a* playing a primary role in modulating innate immunity during adulthood in *C. elegans*.

**Figure 3.**
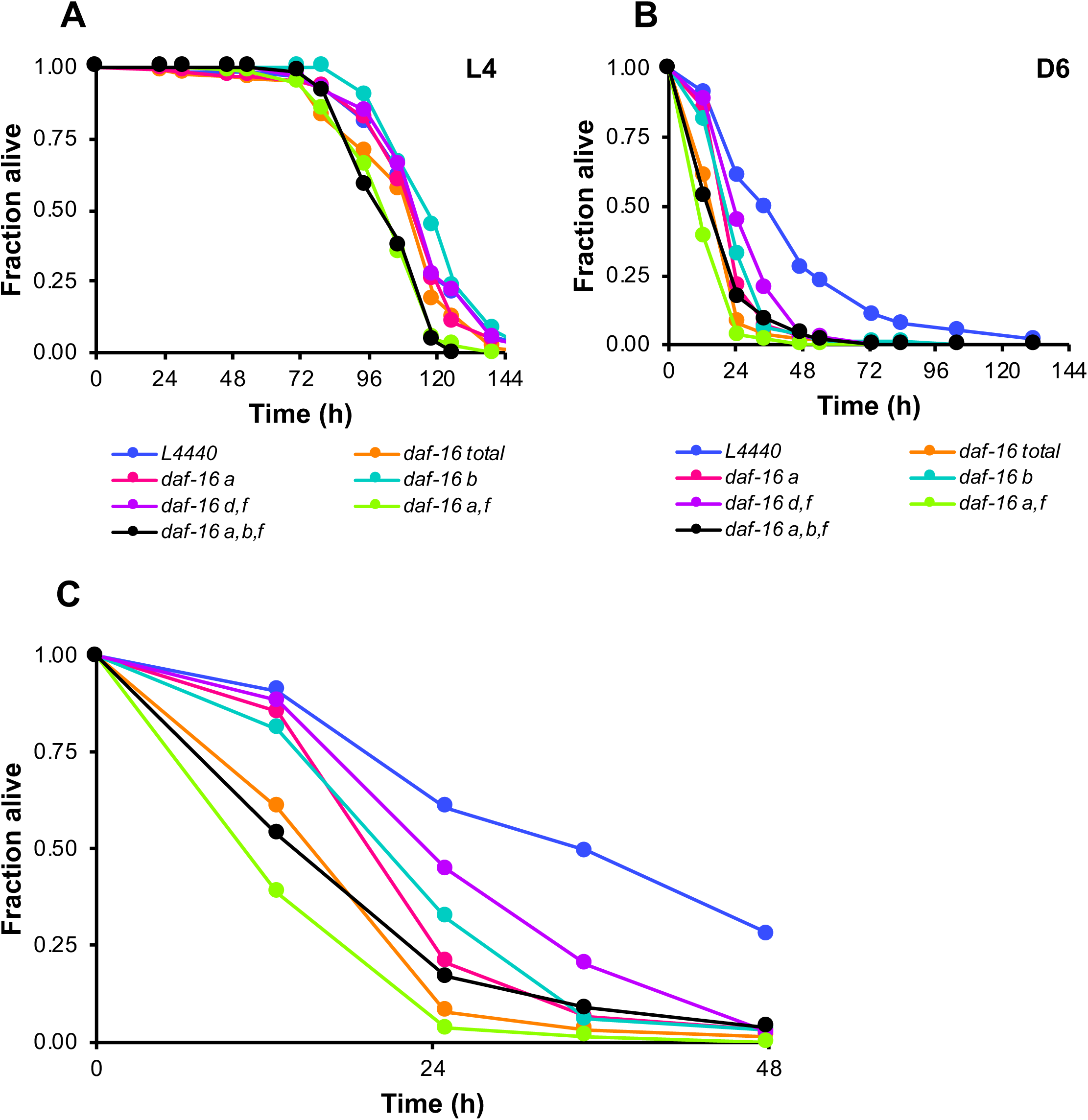
Multiple DAF-16 isoforms contribute to innate immunity in adult *C. elegans*. (A) L4 larvae or (B) Day 6 adults were infected with *P. aeruginosa* PA14 after being treated with RNAi targeting *daf-16* isoforms individually or in combination beginning at the L1 larval stage. L4440 (blue) is an empty RNAi vector control and *daf-16* total (orange) is an RNAi treatment that knocks down all *daf-16* isoforms. Separate RNAi constructs were used to knockdown the following isoforms: *daf-16a* (pink); *daf-16d/f* (purple); *daf-16a*, *b*, and *f* (black); *daf-16b* (turquoise); *daf-16a* and *f* (green). In both panels, the fraction of worms alive is plotted as a function of time. (C) A portion of the survival curve in (B), enlarged to illustrate the effect of knocking down different *daf-16* isoforms. The color scheme is the same as in panels (A) and (B).

### Transcriptional activity of DAF-16 during aging

Considering the homology of DAF-16 to FOXO transcription factors in other species and its well-established function as a transcriptional regulator in *C. elegans*, any influence that it has on innate immunity must be a consequence of the changes in gene expression that it controls. To conduct our own analysis of the transcriptional activity of DAF-16 during aging, we began by first using RNAseq to examine the changes in gene expression that occur between the L4 larval stage and Day 6 of adulthood in wild type N2 animals under the growth conditions that we use during aging experiments in our laboratory to functionally characterize candidate genes. While the expression of most genes changes only modestly between these two time points, we found 1422 genes to be upregulated by 5-fold or greater and 2553 genes to be downregulated by 5-fold or greater (Fig. 4A, Table S1, q<0.05). Among the upregulated genes were those whose products function in the response to cellular stimuli (including stress), regulation of cell death, maintenance of proteostasis, and regulation of signal transduction (Table S2, Fig. S2). Genes downregulated at Day 6 were predominantly associated with metabolic processes, including those that take place in the mitochondria (Table S3, Fig. S3).

**Figure 4.**
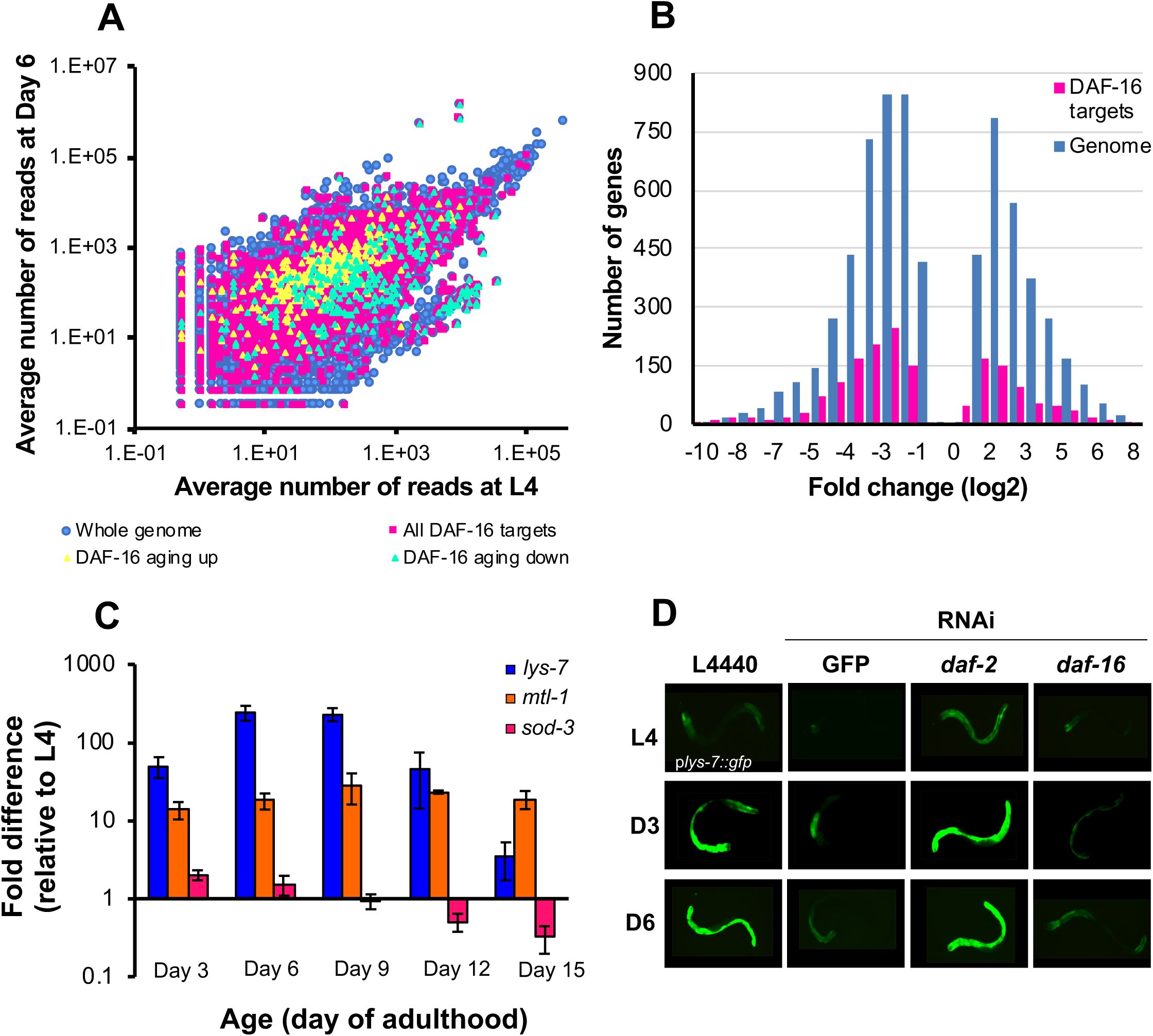
DAF-16 transcriptional activity during aging is dynamic. (A) Differences in gene expression between the L4 larval stage and Day 6 of adulthood are represented in a scatter plot where the average number of reads corresponding to a particular transcript in Day 6 adults is plotted against the average number of reads corresponding to that same transcript in L4 larvae. Each gene is represented as a single dot where blue dots are the entire genome, and pink squares are all DAF-16 targets (see methods for a description for how DAF-16 targets were defined.) Yellow and green triangles are genes reported by Li *et al.* to be up- and downregulated by DAF-16 during aging, respectively. Large changes in gene expression between the two time points lie furthest from the principal diagonal. (B) Histogram showing the distribution of fold changes in gene expression between the L4 larval stage and Day 6 of adulthood in the whole genome (blue) and in DAF-16 targets (pink). (C) Endogenous transcript levels of the DAF-16 targets *lys-7* (blue), *mtl-1* (orange) and *sod-3* (pink) were measured by qRT-PCR at the L4 larval stage and at five points during adulthood. Expression levels were normalized to a housekeeping gene, and the fold difference in gene expression at Days 3, 6, 9, 12, and 15 of adulthood relative to L4 expression levels was calculated by the ΔΔCt method. The average fold change in gene expression at each time point is shown. Error bars represent standard deviation of the mean. (D) The expression of the *in vivo* reporter for DAF-16 activity *plys-7::GFP* was monitored over time by fluorescence microscopy in animals treated with RNAi targeting GFP, *daf-2*, or *daf-16* and compared to control animals treated with an empty RNAi vector, L4440. Shown are representative images of animals at the L4 stage and at Days 3 and 6 of adulthood.

To understand what age-related changes in gene expression might be attributable to DAF-16 transcriptional activity, we created an aggregate catalog of all DAF-16-regulated genes by compiling lists of the transcriptional targets of DAF-16 that have been identified previously (Table S4; McElwee *et al.* 2003; Murphy *et al.* 2003; Tepper *et al.* 2013; Chen *et al.* 2015; Li *et al.* 2019). One set of genes included in our catalog was derived from a recent report that identified genes whose dynamic expression over time requires DAF-16 (Li *et al.* 2019). We designated these as “DAF-16 aging up” or “DAF-16 aging down” to indicate genes that are temporally up- and downregulated by DAF-16, respectively. When the DAF-16 targets in our catalog (3909 genes in total) were mapped onto a scatter plot of gene expression levels at Day 6 versus expression levels at the L4 larval stage, we found that the changes in expression of DAF-16 targets during this period in large part mirror the differential expression of all genes in the *C. elegans* genome between L4 and Day 6 adults (Fig. 4A). Some DAF-16 targets were more highly expressed in adult worms while the expression levels of others declined, suggesting that during aging DAF-16 acts as both a transcriptional activator and repressor. Genes identified as being regulated by DAF-16 during aging partitioned as expected when mapped onto the scatter plot, with “DAF-16 aging up” genes mostly above the principal diagonal and “DAF-16 aging down” genes mostly below it. Overall, the range of the magnitude of the changes in expression between the L4 larval stage and Day 6 of adulthood was similar between DAF-16 targets and the genome as a whole (Fig. 4B). At the same time, DAF-16 seems to be responsible for some of the most dramatic changes in gene expression that take place between L4 and Day 6 of adulthood. We found 348 DAF-16 targets to be upregulated in Day 6 adults (Table S5). Genes encoding products that contribute to the response to stress and DNA replication are overrepresented in this group (Table S6, Fig. S4). 649 DAF-16 targets were detected among genes downregulated between the L4 larval stage and Day 6 (Table S5). Similar to our analysis of genome-wide changes in expression, the expression of DAF-16 transcriptional targets that function in a number of metabolic processes was reduced in post-reproductive adults (Table S7, Fig. S5). Surprisingly, 32 genes purported to function in innate immunity were also overrepresented among the DAF-16 targets that are downregulated in Day 6 animals. In many cases, these genes were assigned to this functional category because they are induced upon exposure to bacteria (Troemel *et al.* 2006; Estes *et al.* 2010). Few have demonstrable roles in immunity, and only six are reported to result in enhanced susceptibility to pathogens when mutated or targeted by RNAi (Wormbase). Our analysis indicates that as animals age DAF-16 actively modulates the expression of at least a subset of its targets, and stress response genes are included among the targets that are upregulated.

While our RNAseq study indicates that DAF-16 transcriptional activity is elevated in Day 6 adults, we wanted to get a better sense of its behavior over the entire course of the worm’s life. Specifically, we wondered when DAF-16 becomes activated during aging and for how long it continues to function as a transcriptional regulator. To answer these questions and to validate the RNAseq results, qRT-PCR was used to measure the age-dependent changes in expression of select DAF-16 targets at the L4 stage and at days 3, 6, 9, 12 and 15 of adulthood (Fig. 4C). We focused our studies on *lys-7*, *sod-3,* and *mtl-1*, three well-established targets of DAF-16 with roles in the response to environmental stressors (Murphy *et al.* 2003). Surprisingly, each of these genes exhibited a unique temporal expression pattern. Relative to their levels in L4 larvae, the expression levels of both *lys-7* and *mtl-1* were elevated throughout adulthood, beginning at least as early as Day 3 and continuing until Day 15. The highest expression of *lys-7* occurred at Day 6 of adulthood gradually declining as worms aged but never dropping to L4 levels of expression. *mtl-1* levels remained relatively constant throughout aging, with little appreciable decline in expression even in the oldest animals tested. The age-dependent expression of *sod-3* followed a completely different pattern. Although there was a small increase in its expression in Day 3 and Day 6 adults, by Day 9 the expression of *sod-3* was slightly less than at L4, and its levels continued to diminish as time went on. Our results indicate that DAF-16 becomes activated in early adults and that it continues to remain transcriptionally active throughout aging, even at very late stages of adulthood.

As an additional line of inquiry into the transcriptional activity of DAF-16 during aging, we sought to uncover *in vivo* evidence for the age-dependent increase in expression of DAF-16 targets using a fluorescent reporter. The most widely used DAF-16 reporter is *sod-3::gfp* in which GFP expression is driven by the promoter of *sod-3*, a gene that encodes a broadly expressed mitochondrial superoxide dismutase (Libina *et al.* 2003). Unlike *lys-7* and *mtl-1*, we did not identify *sod-3* as a gene that is upregulated during adulthood in our RNAseq study, and by qRT-PCR we detected a progressive decrease in its expression in older worms (Fig. 4C). Accordingly, we were unable to detect an increase in *sod-3::gfp* expression during aging by fluorescence microscopy (data not shown). Other *in vivo* sensors of DAF-16 activity were generated in a previous study describing the specificity of the *C. elegans* innate immune response (Alper *et al.* 2007). In one of them the promoter of *lys-7* is fused to *gfp*, driving the expression of GFP in intestinal cells. Since *lys-7* was among the genes that we found to be strongly upregulated throughout adulthood, we chose to study the expression pattern of the *plys-7::gfp* reporter over time after first creating a new strain of worms (VIL001) where *plys-7::gfp* is chromosomally integrated. Microbial infection induces *lys-7* expression, and *lys-7* is required for resistance to bacterial pathogens, suggesting a direct role for LYS-7 in innate immunity as an antimicrobial factor (Mallo *et al.* 2002; Evans *et al.* 2008; Nandakumar and Tan 2008; Kawli and Tan 2008; Boehnisch *et al.* 2011). By homology to the LYS4 gene of the parasite *Entamoeba histolytica*, the product of *lys-7* is predicted to be secreted into the intestinal lumen and to kill bacterial cells through hydrolytic destruction of prokaryotic cell wall peptidoglycans (Leippe 1999; Mallo *et al.* 2002). We monitored p*lys-7::gfp* expression during aging by fluorescence microscopy in a population of age-synchronized animals maintained under standard laboratory conditions with *E. coli* as the food source (Fig 4D, L4440). While the expression of the reporter was initially low in L4 larvae, GFP expression increased by Day 3 of adulthood, and it was robustly expressed in Day 6 animals. To confirm that the age-dependent increase in lys-7::GFP expression was dependent upon DAF-16, we treated animals with RNAi targeting *daf-16* and, as a control, *daf-2*. Whereas knockdown of *daf-2* caused the reporter to be expressed at higher levels than in untreated control animals at all time points (with a rather modest increase in expression at L4), RNAi inhibition of *daf-16* suppressed the age-dependent increase in *plys-7::gfp* expression. These data indicate that *lys-7* is a target of DAF-16 in wild type adult animals, and that DAF-16 is responsible for upregulating *lys-7* during aging. Taken together, our analysis of the transcriptional activity of DAF-16 suggests that during normal aging in unstressed wild type animals a sustained increase in the transcriptional activity of DAF-16 begins during the reproductive phase (by Day 3 of adulthood) and persists throughout adulthood. Our observations indicate that during this time, DAF-16 modulates the expression of a fraction of its complete repertoire of transcriptional targets, some of which contribute to countering challenges from acute stressors.

### Pathogen resistance phenotype of larval and adult *daf-2* mutants

While in younger *C. elegans* DAF-16 is typically inhibited unless and until animals encounter an environmental insult, we found that as wildtype worms age DAF-16 becomes activated even in the absence of an acute stress. One possible mechanism by which DAF-16 becomes activated in adult *C. elegans* is the age-dependent loss of an inhibitory signal through the IIS pathway. We reasoned that if this were the case, then eliminating this inhibition through a loss-of-function mutation in *daf-2* should shift the period in which worms are protected by DAF-16-mediated immunity to begin immediately upon hatching instead of beginning during adulthood. Moreover, we predicted that if derepression of DAF-16 is sufficient to instigate its immune function then the degree of pathogen resistance conferred by disrupting the insulin receptor should be invariant over the worm’s lifespan such that upon infection *daf-2* L4 animals and older post-reproductive *daf-2* worms would experience an equivalent survival advantage relative to their wildtype counterparts. To test our hypothesis, we challenged *daf-2(e1370)* and *daf-2(e1368)* mutants with *P. aeruginosa* at the L4 stage of larval development and at Days 3, 6, and 9 of adulthood. In an allele-specific manner, mutations in *daf-2* conferred varying degrees of resistance to L4 worms (Fig. 5A). Specifically, while a mutation affecting the ligand-binding domain of DAF-2 (*e1368*) did not affect the survival of infected L4 animals, a mutation in the kinase domain (*e1370*) extended their maximum lifespan. Interestingly, we found that the *daf-2* resistance phenotype does not fully manifest until worms become adults. When the *P. aeruginosa* infection was initiated at L4, the survival trajectory of *daf-2(e1370)* mutants paralleled that of wild type worms until ∼96 hours post-infection when their rate of death slowed, causing the median lifespans for all three genotypes in the assay to be similar to each other even though the maximum lifespan of infected *daf-2(e1370)* animals was extended. We found that the resistance phenotype associated with *daf-2* mutations becomes more pronounced in adult animals. When infected at D3 of adulthood the *daf-2(e1368)* mutants that had lacked a phenotype at L4 became resistant to bacterial infection, with longer median and maximum lifespans in the presence of pathogen than wild type animals (Fig. 5B). The *daf-2(e1370)* mutation conferred even greater protection from infection, on average tripling the LT_50_ of D3 adults as compared to wild type. The survival advantage conferred to infected worms by mutations in *daf-2* persisted at Days 6 and 9 of adulthood when wild type animals showed clear signs of immunosenescence (Fig. 5C and D). Corroborating the results of our studies of *daf-2* mutants, RNAi targeting *daf-2* caused Day 6 animals to be more resistant to *P. aeruginosa* infection, but it did not affect the susceptibility of L4 worms (Fig. S6). That inactivation of DAF-2 can extend the lifespan of adult worms infected with a bacterial pathogen suggests that relief of an inhibitory signal through the IIS pathway may be important for activating DAF-16 during adulthood. Yet the fact that the resistance phenotype of *daf-2* mutants can be modified by age raises the possibility that additional factors present in adult worms could also be required for DAF-16-mediated immunity.

**Figure 5.**
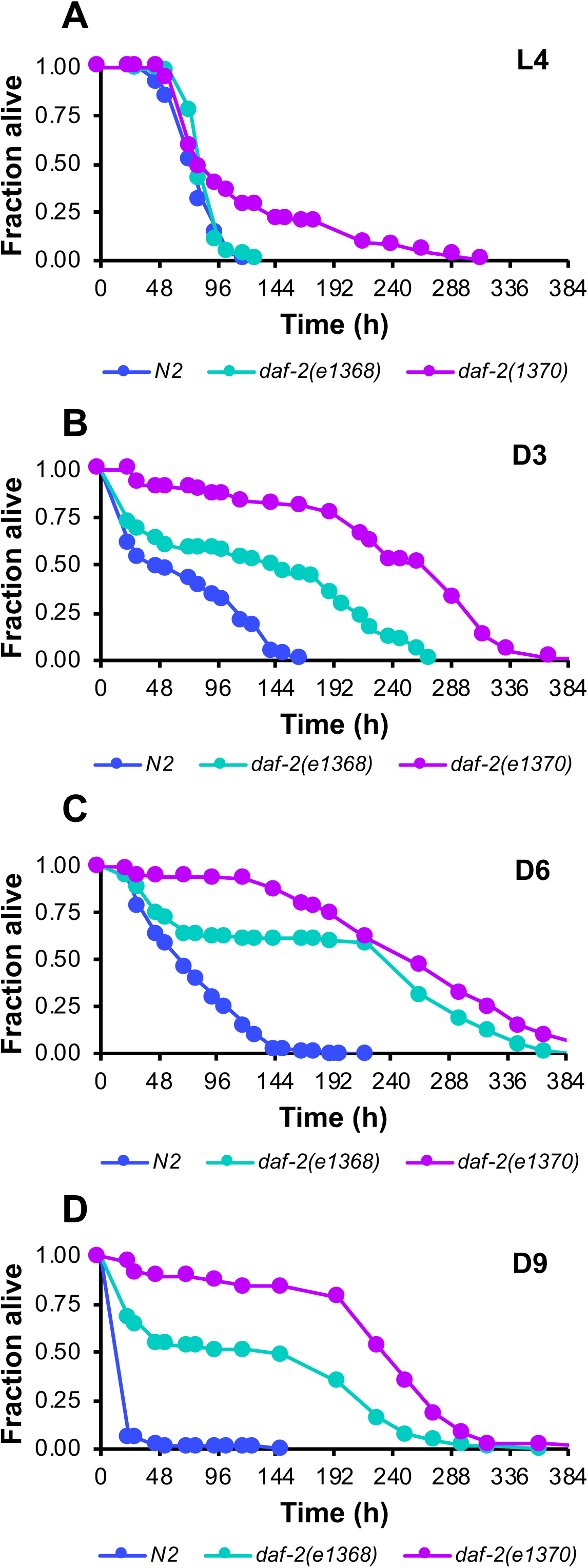
Aging enhances the resistance to bacterial pathogens conferred by *daf-2*. Wild type N2 worms (blue), *daf-2(e1368)* (turquoise), and *daf-2(e1370)* (purple) worms were infected with *P. aeruginosa* at the L4 larval stage (A) or at Day 3, 6, or 9 of adulthood (B-D). Plotted in each panel is the fraction of worms alive versus time.

### Functional analysis of SMK-1 in DAF-16-mediated innate immunity during adulthood

Our data implying a role for DAF-16 in host defense that is restricted to adulthood raises the question of how its activity is regulated as animals age. While the transcriptional activity of DAF-16 is controlled in part through phosphorylation by upstream components of the insulin signaling pathway, its function is also regulated by nuclear factors that control the access of DAF-16 to its targets and influence its transcriptional output (reviewed in Mukhopadhyay *et al.* 2006). Several such factors interact either genetically or physically with DAF-16, yet so far only two, HSF-1 and SMK-1, have been shown to be required for the extended lifespan of *daf-2* mutant (Hsu *et al.* 2003; Samuelson *et al.* 2007). Both are also required for normal lifespan of wildtype animals (Garigan *et al.* 2002; Wolff *et al.* 2006; Lee *et al.* 2009; Chiang *et al.* 2012). HSF-1 and SMK-1 may therefore specifically govern the expression of a subset of DAF-16 targets that promote longevity and survival. Indeed, HSF-1 appears to predominantly control the expression of genes necessary for the response to thermal stress (Hsu *et al.* 2003). SMK-1, in comparison, regulates the expression of DAF-16 targets that are important for counteracting the effects of different stressors including UV radiation, reactive oxygen species, and bacterial pathogens (Wolff *et al.* 2006). We therefore reasoned that SMK-1 may be the more likely candidate to regulate the function of DAF-16 in the host defense of adult *C. elegans*.

To investigate the possibility that SMK-1 regulates DAF-16 in an age-dependent manner, we first asked whether SMK-1 influences the transcriptional activity of DAF-16 in adult worms. RNAi was used to inactivate SMK-1 in a synchronized cohort of worms harboring the *plys-7::GFP* transgene beginning at the L1 larval stage. GFP expression levels were then monitored by fluorescence microscopy at the L4 larval stage and at Day 6 of adulthood (Fig 6A). Mimicking the effect of knocking down *daf-16*, RNAi treatment directed against *smk-1* prevented the age-dependent increase in the expression of the reporter in Day 6 adult worms. This result suggests that SMK-1 is required for the increase in DAF-16 transcriptional activity during aging.

**Figure 6.**
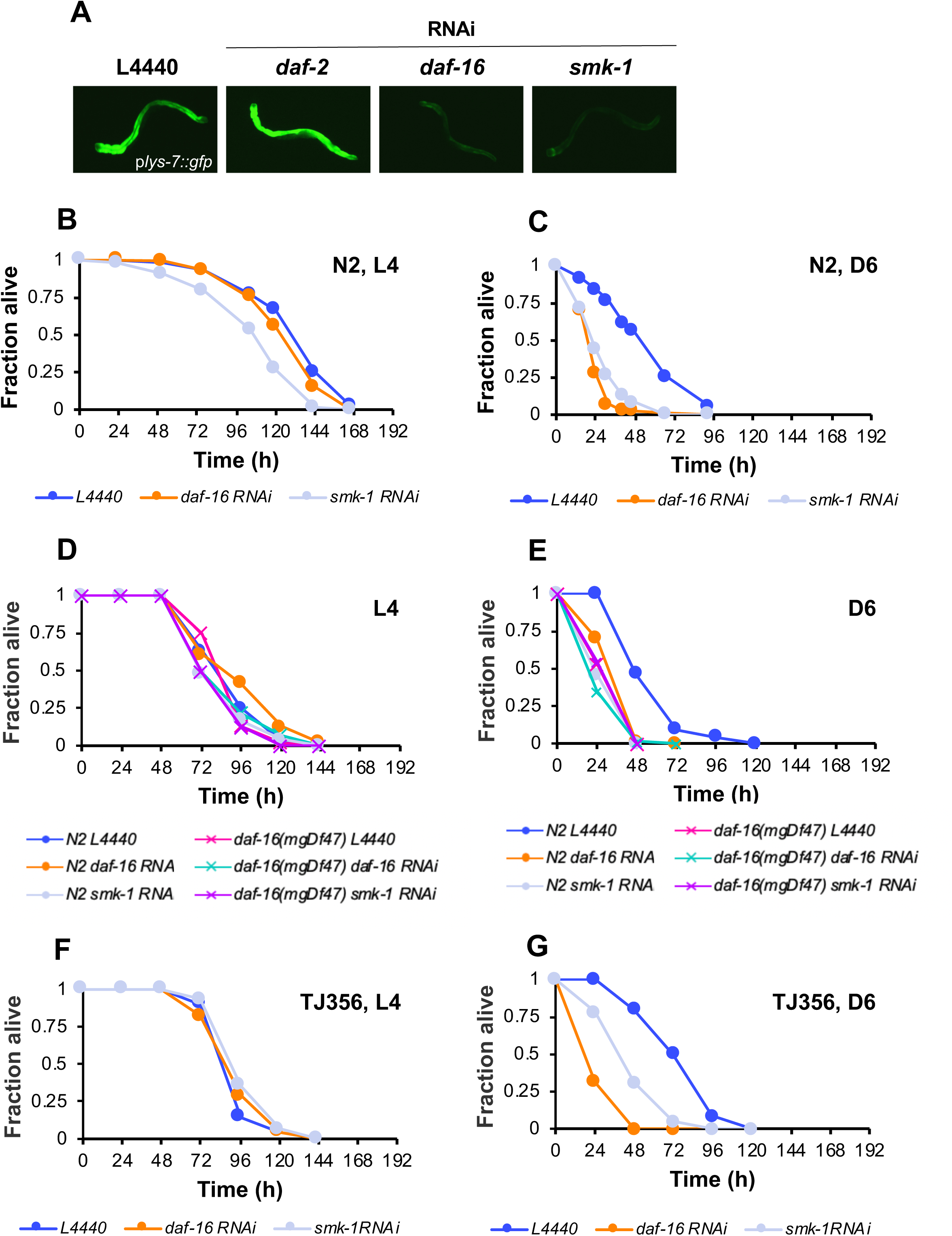
SMK-1 is required for DAF-16-mediated immunity during adulthood. (A) The expression of the DAF-16 reporter *plys-7::GFP* is shown in Day 6 animals that were treated with RNAi targeting the indicated genes beginning at the L1 larval stage. (B) L4 or (C) Day 6 adult N2 wild type worms were infected with *P. aeruginosa* PA14 after being treated with RNAi targeting either *daf-16* (orange) or *smk-1* (grey) beginning at the L1 larval stage. L4440 (blue) is an empty RNAi vector control. (D) L4 or (E) Day 6 N2 wild type worms (filled circles) or *daf-16(mgDf47)* mutants (denoted by X) were infected with *P. aeruginosa* after initiating RNAi treatment to knock down *daf-16* or *smk-1* at the L1 larval stage. The color scheme indicating the RNAi treatments for N2 animals is as described for panels (B) and (C). For *daf-16* mutants pink is the L4440 vector control, turquoise corresponds to *daf-16* RNAi and purple is *smk-1* RNAi. (F) L4 or (G) Day 6 adult TJ356 animals that overexpress DAF-16 were infected with *P. aeruginosa* after being subjected to RNAi to reduce the expression of either *daf-16* (orange) or *smk-1* (grey) beginning at the L1 larval stage. L4440 (blue) is an empty RNAi vector control. The fraction of worms alive is plotted as a function of time.

We next sought to uncover functional evidence to corroborate the results of our transcriptional reporter analysis. Our hypothesis was that if DAF-16-mediated immunity in adult worms is dependent upon SMK-1, then interventions to disrupt the function of SMK-1 should phenocopy the *daf-16* loss-of-function in bacterial infection assays. We again turned to an RNAi approach to explore this possibility. Beginning at the L1 larval stage, *smk-1* was knocked down in a synchronized population of N2 worms, which were then challenged with *P. aeruginosa* at L4 and at Day 6 of adulthood. In a departure from what we observed when inactivating DAF-16 by RNAi, in L4 worms *smk-1* knockdown resulted in a small but significant increase in susceptibility to bacterial pathogen (p= 0.0168, Fig. 6B). Aging exacerbated this phenotype. Similar to the effect of knocking down *daf-16*, RNAi targeting *smk-1* shortened the median survival time of Day 6 adult worms infected with *P. aeruginosa* by almost two-fold compared to untreated control animals (p<0.001, Fig. 6C). These data are consistent with a role for SMK-1 in innate immunity that, similar to DAF-16, becomes more important during adulthood.

That inhibiting the expression of *daf-16* and *smk-1* causes similar phenotypes at Day 6 implies that the two genes act in the same pathway during aging in wild type animals, just as they are reported to do in *daf-2(e1370)* mutants (Wolff *et al.* 2006). If this is the case, then combining a deficiency in both genes in the same animal should not produce an additive effect. To test this prediction and to further investigate the interaction between *smk-1* and *daf-16* during aging, RNAi was used to reduce the expression of *smk-1* in both wild type and *daf-16(mgDf47)* mutants. We then compared their survival to untreated control animals upon infection with *P. aeruginosa* at the L4 larval stage and at Day 6 of adulthood (Fig 6 D, E). Knocking down *smk-1* did not compound the enhanced susceptibility phenotype of *daf-16(mgDf47)* worms when infected as either L4 larvae or as Day 6 adults. That is, at those ages the survival trajectories of *daf-16(mgDf47)* mutants treated with RNAi targeting *smk-1* was similar to that of *daf-16(mgDf47)* control animals and to wild type worms treated with RNAi directed against *daf-16*. These data argue that *daf-16* and *smk-1* function as part of the same genetic pathway during aging in *C. elegans*.

Our functional characterization of *smk-1* during aging indicates that it may be necessary for the increased activity of DAF-16 during adulthood that contributes to host defense. If this is the case, we predicted that *smk-1* would be required to bring about any potential benefit that overexpression of *daf-16* might confer to infected worms. To test this hypothesis, we used RNAi to knock down the expression of *smk-1* or *daf-16* beginning at the L1 larval stage in *C. elegans* strain TJ356 that expresses a DAF-16::GFP fusion from a chromosomally integrated transgene in an otherwise wildtype genetic background (Henderson and Johnson 2001). These animals were then infected with *P. aeruginosa* at either the L4 larval stage or at Day 6 of adulthood. Although *daf-16* overexpression has been reported to improve the survival of L4 animals subjected to thermal stress or UV irradiation (Henderson and Johnson 2001) we did not observe increased resistance to infection of TJ356 worms at the L4 stage (Fig. 6F). The median lifespan of infected L4 TJ356 animals was comparable to that of wild type worms, and it was unchanged by inhibiting the expression of either *daf-16* or *smk-1*. Notably, control TJ356 worms maintained an L4-like median lifespan when they were infected with *P. aeruginosa* at Day 6 of adulthood, suggesting that the effect of overexpressing DAF-16 is to stave off the immunosenescence observed in wild type adults (Fig 6G). This bolstered host defense requires SMK-1. Inhibiting the expression of *smk-1* reduced the median survival of infected Day 6 TJ356 animals, as did knockdown of *daf-16* (Fig 6G). These results indicate that SMK-1 is required for DAF-16-mediated immunity during adulthood.

## DISCUSSION

In this study we sought to determine whether in the face of declining PMK-1 activity in adulthood other pathways might emerge as the predominant mediators of host defense in adult *C. elegans*. For the first time, we demonstrate a functional requirement for DAF-16 in innate immunity specifically during aging. Our functional characterization of DAF-16 over time from the L4 larval stage to the ninth day of adulthood suggests that the timing of its contribution to innate immunity is reciprocal to that of PMK-1. That is, as PMK-1 protein levels fall during aging and therefore minimize its function in immunity, the role of DAF-16 becomes more important after the L4 stage, as it is activated once animals transition to reproductive maturity (Fig. 1). Our results suggest that the transcriptional activity of DAF-16 increases in an age-dependent manner, even in wild type worms maintained under normal conditions in the absence of any environmental stresses (Fig. 4). Both the transcriptional activity of DAF-16 and its role in immunity in adult worms require *smk-1* (Fig. 6), suggesting that, as is the case when it is activated in response to stress in younger worms, DAF-16 cooperates with other proteins to carry out its function in adults (Mukhopadhyay *et al.* 2006). Taken together, the observations we report here indicate that not all pathways that protect animals from infection are in decline during aging. Instead, in adulthood immunity is remodeled such that there may be a shift from one set of immune pathways that act primarily earlier in life to other pathways that take over the duty of host defense later on.

The contribution of DAF-16 to lifespan was established in studies demonstrating that it is required for the extended lifespan of *daf-2* mutants (Kenyon *et al.* 1993). The critical temporal window for this activity of DAF-16 appears to occur not during larval development but since delaying knockdown of daf-16 in *daf-2(e1370)* mutants until early adulthood is sufficient to completely suppress the *daf-2* longevity phenotype (Dillin *et al.* 2002). Waiting even longer to block the function of DAF-16 (until Day 6 or even Day 15 of adulthood) by RNAi treatment still partially suppresses *daf-2*, shortening the lifespan of *daf-2(e1370)* to some degree. These experiments demonstrate that DAF-16 is capable of functioning even late into adulthood in *C. elegans*, especially under conditions in which signaling through the IIS pathway is disrupted. Whether DAF-16 is normally activated as part of the aging process in wild type animals, however, cannot be discerned from these studies.

Since DAF-16 must translocate from the cytosol to the nucleus to carry out its cellular function as a transcriptional regulator, more direct assessments of the timing of DAF-16 activation during aging have largely relied upon detecting and in many cases quantifying the proportion of nuclear localized DAF-16 over the course of the worm’s life, with the caveat that the mere presence of DAF-16 in the nucleus is not sufficient for its activation (Henderson *et al.* 2006; Berdichevsky *et al.* 2006). Interestingly, even in cases where DAF-16 is found to be predominantly cytosolic, its transcriptional targets are still expressed (Dues *et al.* 2016) and physiological processes that are dependent upon DAF-16 activation appear to go on as normal (Baxi *et al.* 2017). These observations make evident the need for functional characterization to substantiate conclusions regarding the activity of DAF-16 based on its subcellular localization. Our studies address this deficiency by systematically testing the functional requirement for DAF-16 over time, and they establish a role for DAF-16 in innate immunity that is exclusive to adulthood.

Despite indications that DAF-16 activity may be repressed by fertilized oocytes during reproduction (Miyata *et al.* 2008), a majority of studies of DAF-16 localization have found that at least a fraction of DAF-16 is nuclear early on in reproduction (Lin *et al.* 2001; Weinkove *et al.* 2006; Tepper *et al.* 2013; Baxi *et al.* 2017; Li *et al.* 2019), and functional evidence verifies that it is active at that time (Baxi *et al.* 2017; Li *et al.* 2019). What happens later, however, is somewhat controversial. Some studies show an efflux of DAF-16 out of the nucleus as worms age with a corresponding decline in efficiency of DAF-16-dependent functions that becomes evident at the end of reproduction (Weinkove *et al.* 2006; Tepper *et al.* 2013; Baxi *et al.* 2017). Others, however, point to the beginning of the post-reproductive period as the very time when DAF-16 nuclear enrichment is at a maximum (Gurkar *et al.* 2018; Li *et al.* 2019). This is coincident with the point at which an increase in the expression of the *daf-16a* and *daf-16d/f* isoforms that becomes most pronounced after the L4 stage reaches its peak (Bansal *et al.* 2014). Consistent with a recent report demonstrating increased expression of DAF-16 targets at Day 2-3 of adulthood (Li *et al.* 2019), we were able to narrow the timing of the onset of increased DAF-16 activity during aging to within the first three days of adulthood in part because we detected elevated levels of genes regulated by DAF-16 as early as Day 3 of adulthood (Fig. 4C). The expression of some of these genes increased even further as worms aged and remained elevated for more than half of the worm’s lifespan, suggesting that there may be a progressive increase in the transcriptional activity of DAF-16 over time. This is likely directed particularly at select genes, as only a subset of the total catalog of DAF-16 targets were differentially regulated during aging (Fig. 4A, Table S5). Fine tuning of the transcriptional output of DAF-16 in adult animals may be achieved at least in part by differences in the relative activity of DAF-16 isoforms that were recently shown to have both discrete and overlapping targets (Chen *et al.* 2015). Our functional analysis of *daf-16* isoforms supports this possibility (Fig. 3).

While the subset of transcriptional targets of DAF-16 that are upregulated during aging encodes products that contribute to a number of different cellular processes (Table S6), we found that there is a discrete functional requirement for DAF-16 in innate immunity in adult *C. elegans*. Neither *daf-16* loss-of function mutants nor animals treated with RNAi targeting *daf-16* were more susceptible to *P. aeruginosa* at the L4 larval stage, but they became more susceptible to the pathogen when infected as adults (Figs. 1 and 2). This is the opposite of the scenario that was previously reported for *pmk-1(km25)* mutants (Youngman *et al.* 2011) and that we recapitulated here where *pmk-1(km25)* larvae exhibit a more severe sensitivity to pathogen than adults (Fig. 1). Therefore, whereas PMK-1 seems to contribute to innate immunity in larvae and younger adults, in a reciprocal fashion, DAF-16 becomes increasingly important for host defense as animals age. In fact, our data suggest that it is required exclusively during adulthood to confer resistance to bacterial pathogens (Fig. 2E-G), mirroring its timing requirement for lifespan determination (Dillin *et al.* 2002).

The activation of DAF-16 during aging does not depend upon an induced response to environmental insults, which in any case appears to wane during adulthood (Dues *et al.* 2016). Rather, we observed an age-dependent increase in DAF-16 activity in the absence of acute stress (Fig 4C, D). Although the identity of the upstream trigger that stimulates DAF-16 to become activated during aging is not known, one possibility is that the repressive signal through the IIS pathway may be attenuated as animals age. In our hands animals overexpressing DAF-16 died from infection at the same rate as control animals when challenged at the L4 larval stage, but they became resistant to bacterial infection at Day 6 of adulthood (Fig. 6F, G). If all of the additional copies of DAF-16 present in the transgenic animals were immediately activated then the L4 animals should have also been resistant to *P. aeruginosa* infection because they would have been expressing higher levels of immune effectors leading up to their encounter with the pathogen. That the benefit of DAF-16 overexpression is delayed until adulthood is consistent with a model in which inhibition of DAF-16 via the IIS pathway is counteracted in an age-dependent manner. Consistent with this possibility, the pathogen resistance phenotype associated with the two hypomorphic *daf-2* mutations that we studied (*e1368* and *e1370*) did not fully manifest until worms reached adulthood (Fig. 5). Putative negative regulation of the IIS pathway with age is likely to occur downstream of the DAF-2 receptor since the transcript levels of its agonists do not decline over time (Li *et al.* 2019).

Regardless of the nature of the upstream input that stimulates DAF-16 activation in adult worms, our data indicate that its age-dependent function requires SMK-1. SMK-1 is the *C. elegans* ortholog of an evolutionarily conserved regulatory subunit of the phosphoprotein phosphatase 4 (PP4) complex (Kim *et al.* 2007). In diverse species, the PP4 catalytic subunit forms a trimeric complex with two other proteins that regulate its enzymatic function and determine its substrate specificity (Cohen *et al.* 2005; Sousa-Nunes *et al.* 2009). Acting to antagonize the function of a host of kinases, the PP4 complex carries out critical regulatory functions. For example, in vertebrates PP4 is important for the proliferation and homeostatic expansion of T cells (Liao *et al.* 2014), and it positively regulates VDJ recombination during B cell development (Su *et al.* 2013). In both yeast and mammals, PP4 functions in the response to genotoxic stress by limiting the duration of DNA damage checkpoints and promoting the recovery from double strand break repair (Lee *et al.* 2010, 2012; Kim *et al.* 2011; Shaltiel *et al.* 2014). The *C. elegans* ortholog of the catalytic subunit of the PP4 complex is PPH-4.1, and in an *in vitro* expression system it physically interacts with SMK-1 (Kim *et al.* 2007). During embryogenesis in *C. elegans*, the SMK-1/PPH-4.1 complex functions to establish asynchrony in the progenitors of different cell lineages by preventing the ATR-Chk1 checkpoint pathway from responding to DNA damage. As a result, ATR-Chk1 controls the timing of cell divisions exclusively in response to developmental cues.

Separate from its role in development, SMK-1 plays a role in lifespan determination in *C. elegans*, as evidenced primarily through functional characterization of SMK-1 in *daf-2(e1370)* mutants (Wolff *et al.* 2006). For example, the extended lifespan of *daf-2(e1370)* mutants requires *smk-1*. Similar to knockdown of *daf-16*, inhibiting the expression of *smk-1* suppresses the resistance of *daf-2(e1370)* mutants to oxidative stress, ultraviolet irradiation, and bacterial infection. At the molecular level the increased level of expression of several DAF-16 transcriptional targets in *daf-2(e1370)* animals, including *sod-3*, is dependent upon *smk-1*. These data indicate that SMK-1 modulates the function of DAF-16 under conditions in which DAF-16 is constitutively activated because of the absence of an inhibitory signal through the IIS pathway. Our work builds upon these observations. We found that SMK-1 contributes to longevity not only in genetic backgrounds where DAF-16 is chronically derepressed but also during normal aging in wildtype animals by facilitating DAF-16-mediated stress resistance. Similar to *daf-16* loss-of-function mutants, animals subjected to RNAi targeting *smk-1* show a more pronounced sensitivity to bacterial infection as adults than they do as late larvae (Fig. 6B, C), suggesting that the functional requirement for SMK-1 is predominantly during adulthood as it is for DAF-16.

In light of our functional characterization of SMK-1 in adult worms and our observation that the upregulation of an *in vivo* reporter of DAF-16 during aging requires SMK-1 (Fig. 6), we propose a model in which SMK-1 is necessary for the age-dependent transcriptional regulatory function of DAF-16 (Fig. 7). While DAF-16 and SMK-1 are coexpressed in the nucleus of several tissues in *C. elegans* (Wolff *et al.* 2006), there is no evidence of a direct or indirect physical interaction between the two proteins. Therefore, our model instead invokes biochemical evidence showing SMK-1 to associate with PPH-4.1 (Kim *et al.* 2007). SMK-1 would thus exert its influence over DAF-16 as part of the *C. elegans* PP4 complex by regulating the activity of the PPH-4.1 phosphatase (Kim *et al.* 2007). While substrates of the PP4 complex in worms have not yet been identified, it is tempting to speculate that, similar to its function in DNA repair in other organisms, during aging the PPH-4.1/SMK-1 PP4 complex reverses repressive marks on histones at key genomic loci to allow for access to the promoters of genes that contribute to homeostasis, including defense, detoxification, repair, and maintenance. In this way, SMK-1 would modulate both the activity and the transcriptional output of DAF-16 through an epigenetic mechanism reminiscent of PP4 regulation of Smad1 target expression through the inhibition of HDAC3 in zebrafish (Jia *et al.* 2012). As the upregulation of DAF-16 targets in *daf-2(e1370)* mutants depends upon the SWI/SNF complex (Riedel *et al.* 2013) and considering the widespread reorganization of chromatin during aging (Feser and Tyler 2011), it seems plausible that DAF-16 transcriptional activity in adult animals would also require manipulation of chromatin structure.

**Figure 7.**
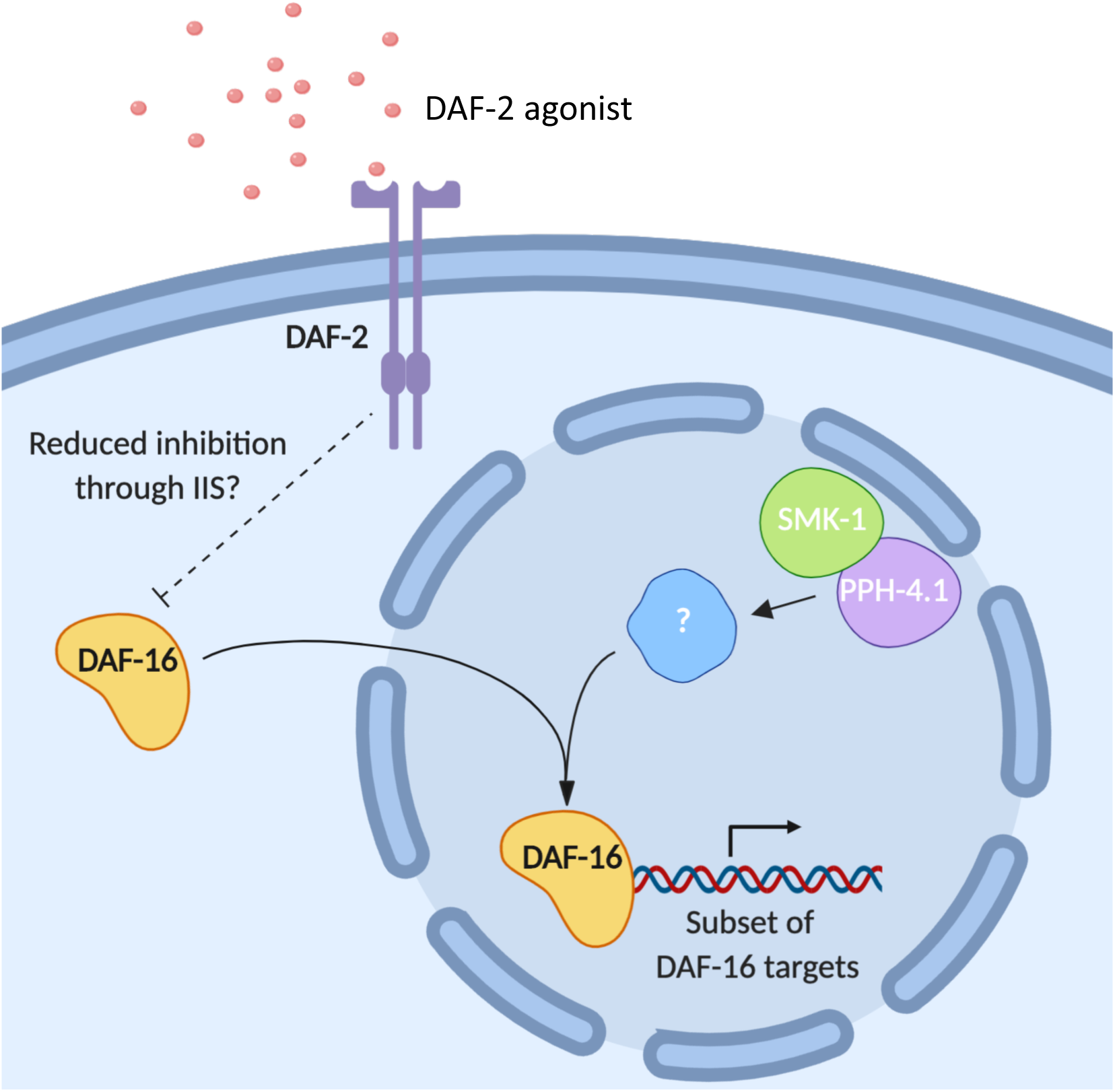
DAF-16 and SMK-1 cooperate to modulate innate immunity during aging. During early adulthood, DAF-16 is permitted to enter the nucleus where it upregulates a subset of its transcriptional targets, including immune effectors. The age-dependent increase in DAF-16 transcriptional activity appears to require SMK-1, a regulatory subunit of the *C. elegans* phosphoprotein phosphatase 4 (PP4) complex whose substrates in the context of aging have not yet been identified. Since SMK-1 localizes to the nucleus, we speculate that the substrate of the SMK-1/PPH-4.1 complex that influences the transcriptional activity of DAF-16 is also nuclear.

The shift to predominantly DAF-16-mediated immunity during adulthood may be a matter of energetic economy. Age-related changes to feeding behavior in worms reduces the intake of food (Huang *et al.* 2004), potentially creating a scarcity of resources to be distributed among multiple layers of host defense. In anticipation of challenges that are yet to be encountered later in life, devoting what resources remain to activating molecules like DAF-16 that confer protection from a broad spectrum of environmental insults would allow the worm to diversify its means of responding to stressors in return for a relatively small energetic investment.

## ACKNOWLEDGEMENTS

We thank Dennis Kim and members of his laboratory for their generous gifts of resources, reagents, and advice. Portions of this work were funded by a grant to MJY from the Charles E. Kaufman Foundation (#KA2014-73918). DRM was funded in part by a teaching assistantship through the Villanova Department of Biology. PR and PP received undergraduate research fellowships from the Villanova Center for Undergraduate Research and Fellowships. SB was awarded a Mary E. Desmond Fellowship from the Villanova University Department of Biology.

We acknowledge the office of the Dean of the College of Liberal Arts and Sciences at Villanova for additional funding. Some strains were provided by the CGC, which is funded by NIH Office of Research Infrastructure Programs (P40 OD010440). We are grateful to Walter Fontana’s laboratory for sharing the original *plys-7::gfp* reporter strain.

---

**Figure S1. Second replicate of RNAi targeting *daf-16* isoforms.** (A) L4 larvae or (B) Day 6 adults were infected with *P.aeruginosa* PA14 after being treated with RNAi targeting *daf-16* isoforms individually or in combination beginning at the L1 larval stage. L4440 (blue) is an empty RNAi vector control and *daf-16* total (orange) is an RNAi treatment that knocks down all *daf-16* isoforms. Separate RNAi constructs were used to knockdown the following isoforms: *daf-16a* (pink); *daf-16d/f* (purple); *daf-16a*, *b*, and *f* (black); *daf-16b* (turquoise); *daf-16a* and *f* (green). In both panels, the fraction of worms alive is plotted as a function of time. (C) A portion of the survival curve in (B), enlarged to illustrate the effect of knocking down different *daf-16* isoforms. The color scheme is the same as in panels (A) and (B).

**Figure S2. Gene ontology classifications for genes upregulated by at least 5-fold between the L4 larval stage and Day 6 of adulthood.** Pie charts represent GO biological process terms (A) or GO cellular process terms (B).

**Figure S3. Gene ontology classifications for genes downregulated by at least 5-fold between the L4 larval stage and Day 6 of adulthood.** Pie charts represent GO biological process terms (A) or GO cellular process terms (B).

**Figure S4. Gene ontology classifications for DAF-16 targets upregulated by at least 5-fold between the L4 larval stage and Day 6 of adulthood.** Pie charts represent GO biological process terms (A) or GO cellular process terms

**Figure S5. Gene ontology classifications DAF-16 targets downregulated by at least 5-fold between the L4 larval stage and Day 6 of adulthood.** Pie charts represent GO biological process terms (A) or GO cellular process terms (B).

**Figure S6. Manipulation of the IIS pathway predominantly affects adult and not larval stage immunity.** The survival of wild type N2 worms (blue) was compared to strains bearing loss-of-function mutations affecting the insulin receptor (*daf-2(e1368)*, turquoise, or *daf-2(e1370),* purple) or the DAF-16 transcription factor (*daf-16(mgDf47)*, orange) upon infection with *P. aeruginosa* at the L4 larval stage (A) or Day 3 (B) or Day 6 (C) of adulthood. After initiating RNAi treatment to knockdown either *daf-2* (turquoise) or *daf-16* (orange) at the L1 stage, N2 wild type animals were infected with *P. aeruginosa* as L4 larvae (D) or as Day 6 adults (E). L4440 is an empty RNAi vector control (blue). In each panel the fraction of animals alive is plotted as a function of time.

